# Dynamin is required for efficient cytomegalovirus maturation and envelopment

**DOI:** 10.1101/396820

**Authors:** Mohammad H. Hasan, Leslie E. Davis, Ratna K. Bollavarapu, Dipanwita Mitra, Rinkuben Parmar, Ritesh Tandon

## Abstract

Cytomegalovirus secondary envelopment occurs in a virus-induced cytoplasmic assembly compartment (vAC) generated via a drastic reorganization of the membranes of the secretory and endocytic systems. Dynamin is a eukaryotic GTPase that is implicated in membrane remodeling and endocytic membrane fission events; however, the role of dynamin in cellular trafficking of viruses beyond virus entry is only partially understood. Mouse embryonic fibroblasts (MEF) engineered to excise all three isoforms of dynamin were infected with mouse cytomegalovirus (MCMV-K181). Immediate early (IE1; m123) viral protein was detected in these triple dynamin knockout (TKO) cells as well as in mock-induced parental MEF at early times post infection although levels were reduced in TKO cells, indicating that virus entry was affected but not eliminated. Levels of IE1 protein and another viral early protein (m04) were normalized by 48 hours post infection; however, late protein (m55; gB) expression was significantly reduced in infected TKO cells compared to parental MEF. Ultrastructural analysis revealed intact stages of nuclear virus maturation in both cases with equivalent numbers of nucleocapsids containing packaged viral DNA (C-capsids) indicating successful viral DNA replication, capsid assembly and genome packaging. Most importantly, severe defects in virus envelopment were visualized in TKO cells but not in parental cells. Dynamin inhibitor (dynasore) treated MEF showed a phenotype similar to TKO cells upon MCMV infection confirming the role of dynamin in late maturation processes. In summary, dynamin-mediated endocytic pathways are critical for the completion of cytoplasmic stages of cytomegalovirus maturation.

**Importance:** Viruses are known to exploit specific cellular functions at different stages of their life cycle in order to replicate, avoid immune recognition by the host and to establish a successful infection. Cytomegalovirus (CMV) infected cells are characterized by a prominent cytoplasmic inclusion (virus assembly compartment; vAC) that is the site of virus maturation and envelopment. While endocytic membranes are known to be the functional components of vAC, knowledge of specific endocytic pathways implicated in CMV maturation and envelopment is lacking. Here we show that dynamin, which is an integral part of host endocytic machinery, is largely dispensable for early stages of CMV infection but is required at a late stage of CMV maturation. Studies on dynamin function in CMV infection will help us understand the host-virus interaction pathways amenable to targeting by conventional small molecules as well as by newer generation nucleotide-based therapeutics (e.g. siRNA, CRISPR/CAS gRNA, etc.).

## Introduction

Endocytic pathways are important for cellular entry of several viruses (1-5); however, their role in post-entry stages of virus replication is far from resolved. Maturing herpesvirus nucleocapsids undergo primary envelopment at the inner nuclear membrane, traverse through the nuclear envelope, uncoat at the outer nuclear membrane and reach the cytoplasm where secondary or final envelopment takes place (6, 7). The cytoplasmic stage of herpesvirus maturation has been particularly challenging to study because a myriad of host and viral factors contribute to this process (6, 8). The identity of the cellular membranes that contribute to final virus envelope has been a topic of several studies (9-11). A further challenge in these studies has been the possibility of mislocalization of cellular markers during infection. To elaborate this point, biomarkers that associate with the endoplasmic reticulum (ER) may not associate with the ER during infection or the ER membranes may form completely different structures during infection. For human cytomegalovirus (HCMV), elegant 3-dimensional confocal studies have shown the organization of a virus assembly compartment (vAC) in the cytoplasm, the site of virus maturation (12, 13). The vAC consists of several host organelles organized in specific shape and capacity with early endosomes forming the core of the structure. A similar vAC has been described for mouse cytomegalovirus (MCMV) infected cells (14). Moreover, endosomal processes have been implicated in cytoplasmic maturation of several herpesviruses (15-19). Endocytic motifs in herpes simplex virus (HSV) envelope glycoprotein B (gB) are required for proper recycling of gB from cell surface to trans-Golgi network during maturation and thereby determine the infectivity of maturing virus (20). Similar endocytic processes for internalization of pseudorabies virus and CMV gB have also been reported (21, 22) and there is evidence that endocytic membranes are used for envelopment of several herpesviruses including HSV, varicella-zoster virus (VZV), and CMV (10, 13, 17, 23, 24).

Dynamins and dynamin-related proteins (DRP) constitute a superfamily of large self-assembling GTPases (an enzyme that can bind and hydrolyze guanosine triphosphate (GTP)) that mediate membrane fission and fusion in biological processes such as endocytosis, vesicle trafficking, cell division, organelle division and fusion (25). They are distinct from the small, Ras-like GTPases due to their oligomerization-dependent activation, the capacity to interact directly with membrane lipids, and their low GTP binding activity (26). Dynamins work twice in the mechanism of endocytosis: early in the constriction of the invaginating vesicle and late in its scission (27). Dynamins are known to be required for clathrin-mediated endocytosis (28, 29). In mammals, classical dynamins include dynamins 1, 2 and 3. Dynamin 1 is enriched within the brain and localizes to presynaptic terminals, dynamin 2 has a ubiquitous tissue distribution, whereas dynamin 3 is localized in the testis and the brain (25).

Earlier, we studied the process of HCMV maturation in cells where dynamin-clathrin pathways were pharmacologically inhibited (30). One of the small molecules, dynasore, used in this study specifically inhibits dynamin function (31). In the current study, we utilized the recently established conditional triple dynamin knockout mouse embryonic fibroblasts (TKO) (32) to study the involvement of endocytic pathways in cytomegalovirus maturation. The use of TKO cells over the drug is preferred because adverse side effects of the dynamin inhibitor dynasore cannot be entirely ruled out. The results of this study reveal that dynamin is critical for a late stage of virus maturation. Studies on dynamin function in herpesvirus infection will help us understand the host-virus interaction pathways amenable to targeting by conventional small molecules as well as by newer generation nucleotide-based therapeutics (e.g. siRNA, CRISPR/Cas9 gRNA, etc.). Targeting dynamin with pharmaceutical compounds has already been shown to have prophylactic potential against several infectious agents (reviewed in (33)).

## Materials and Methods

### Cells

Mouse embryonic fibroblasts (MEF) were cultured in Dulbecco’s modified Eagle’s medium (DMEM, Cellgro, Manassas, VA) containing 4.5 g/ml glucose, 10% fetal bovine serum SAFC, Lenexa, KS), 1 mM sodium pyruvate, 2 mM L-glutamine, and 100 U/ml penicillin-streptomycin (Cellgro, Manassas, VA) at 37°C with 5% CO_2_. The deletion of dynamin in engineered triple dynamin knockout (TKO) cells is mediated by a tamoxifen inducible knockout strategy. Briefly, these cells express a Cre-estrogen receptor mutant knock-in transgene from the ROSA26 locus (34). Thus, Cre is only shuttled into the nucleus in response to tamoxifen exposure. The TKO cells were treated with 10 µM stock of 4-hydroxytamoxifen (4-HT, Sigma H-6278) in 100% ethyl alcohol for 2 days and then media was changed back to normal tamoxifen-free media. Depletion of dynamin was evident at 3-4 days post treatment in western blots (described below) of whole cell lysates.

### Antibodies, immunofluorescence assays, and immunoblots

The mouse anti-dynamin clone 41 from BD (#610245) was used to probe for dynamin in the Western blotting. Mouse cytomegalovirus IE1 (m123), m04, m06 and m55 mouse antibodies (Catalog nos. HR-MCMV-08, HR-MCMV-01, HR-MCMV-02 and HR-MCMV-04) were purchased from Center for Proteomics, University of Rijeka, and used at 1:1000 dilution. Golgin-97 rabbit antibody was purchased from Cell Signaling Technology (Catalog No. 13192S) and used at 1:1000 dilution. Fluorescent label tagged secondary antibody DYLIGHT 594 was purchased from Thermo Scientific Pierce and used at 1:1000 in immunofluorescent assays (IFA) described below. Hoechst 33258 (Thermo Scientific Pierce) staining (1:3000 dilution) identified the nuclei in IFA. Anti β-actin antibody (AC-74, Sigma-Aldrich, St. Louis, MO) was used (1:1000 dilution) as a control for sample loading in immunoblots (IB). Horseradish peroxidase-labeled anti-mouse IgG, IgM and anti-Rabbit IgG (Catalog Nos. 31444 and 31460, Thermo Scientific, Rockford, IL) were used as the secondary antibody at 1:3000 dilutions for IBs. Blots were detected using ECL Western blotting detection reagents (GE Healthcare, Buckinghamshire, United Kingdom).

### Virus

MCMV strain K181 was grown in MEF cells. Virus stock was prepared in 3X autoclaved milk, sonicated 3 times and stored at −80°C. During infection, media was removed from the wells of cell culture plates and appropriately diluted virus stock was absorbed onto the cells in raw DMEM. Cells were incubated for 1 hour with gentle shaking every 10 mins followed by washing 3X with PBS. Fresh complete medium was added and cells were incubated until the end point.

### Cell Viability Assay

Parental MEF and TKO cells grown on 12 well tissue culture plates were infected with MCMV-K181 at a multiplicity of infection (MOI) of 3.0 or mock-infected at confluency. Five hundred µl of fresh complete medium was added to the wells on day 3 and day 6. At the designated time points, media was removed and cells were harvested by trypsinization. Cell viability was determined using trypan blue exclusion on TC20 automated cell counter (BioRad Laboratories, Hercules, CA) following manufacturer’s protocol.

### Microscopy

Samples were prepared using established protocols for IFA and confocal fluorescence microscopy. Briefly, mock induced parental MEF or 4-HT treated TKO cells were grown on coverslip-inserts in 24 well tissue culture dishes and infected with an MOI of 3.0 at confluency. At the end point of the experiment, cells were fixed in 3.7% formaldehyde for 10 min and were incubated in 50 mM NH_4_Cl in 1X PBS for 10 min to reduce autofluorescence. This was followed by washing in 1X PBS, incubation in 0.5% Triton X-100 for 20 min to permeabilize the cells and finally washing and incubation with primary and secondary antibodies at 1:1000 dilution in 0.1% bovine serum albumin in 1X PBS. Coverslips were retrieved from the wells and were mounted on glass slides with a drop of mounting medium (Gel/Mount, Biomeda, Foster City, CA) and dried overnight before imaging. Images were acquired on an inverted Evos-FL microscope (Thermo Fisher Scientific, Waltham, MA) using 100X objective. Samples for transmission electron microscopy (TEM) were prepared by fixing the cells (MEF) at endpoint in 2.5% glutaraldehyde in 0.1 M cacodylate buffer (pH 7.2) for 2 h at room temperature. Cells were then washed with the same buffer and postfixed with buffered 1.0% osmium tetroxide at room temperature for 1 h. Following several washes with 0.1M cacodylate buffer, cells were dehydrated with ethanol, infiltrated, and embedded in Eponate 12 resin (Ted Pella Inc., Redding, CA). Cell culture plates were cracked with a hammer to release the resin after it had solidified, and ultrathin sections (60 to 70 nm) of monolayer cells were cut and counterstained using uranyl acetate and lead citrate. Examination of ultrathin sections was carried out on a Hitachi H-7500 TEM operated at 75 kV, and images were captured using a Gatan BioScan (Pleasanton, CA) charge-coupled device camera. The images were acquired and analyzed with the Digital Micrograph (Pleasanton, CA) software.

### Drug inhibition assay

Confluent MEF monolayers were pretreated with dynasore (50 µM) (Catalog No. 324410, EMD Millipore Corp., Billerica, MA) for 1 h and infected with MCMV-K181 in medium containing the drug. Cells were washed with PBS and then incubated in the presence of the drug until the end of the experiment.

### Virus titers

Infected or mock-infected samples were harvested within the medium at the designated end points and stored at −80°C before titration. In some experiments, media and cells were separated by low-speed (< 1000 X g) centrifugation and viral loads in supernatant and cells were quantified by titering on wild-type MEF. Titers was performed as described earlier (35) with some modifications. In brief, monolayers of MEF grown in 12 well plates and serial dilutions of sonicated samples were absorbed onto them for 1 h, followed by 3X washing with PBS. Carboxymethylcellulose (CMC) (Catalog No. 217274, EMD Millipore Corp., Billerica, MA) overlay with complete DMEM media (1-part autoclaved CMC and 3 parts media) was added and cells were incubated for 5 days. At end point, overlay was removed and cells were washed 2X with PBS. Infected monolayers were fixed in 100% methanol for 7 min, washed once with PBS and stained with 1% crystal violet (Catalog No. C581-25, Fisher Chemicals, Fair Lawn, NJ) for 15 min. Plates were finally washed with tap water, air dried and plaques with clear zone were quantified.

## Results

### MCMV replicates to low titers in dynamin-depleted fibroblasts

Dynamin was depleted in engineered MEF by treatment with 4-HT for 48 hours (Fig 1A) leading to the generation of TKO cells as described before (32). Whole cell lysates of MCMV-K181 infected (+) or mock-infected (-) MEF showed similar levels of dynamin whereas dynamin was reduced to insignificant levels in TKO cells. Mock-induced parental MEF and TKO cells were infected with MCMV-K181 at a high (3.0) or low (0.01) multiplicity of infection (MOI) and monitored for virus growth. Cells were harvested in the medium at 3- or 6-days post infection and analyzed for plaque forming units on wild-type MEFs. At 3 days post infection, MCMV titers were reduced about 10-fold for both low and high MOI infections in TKO cells compared to parental MEF (Fig 1B). At 6 days post infection, TKO virus titers were reduced more than 200-fold for low MOI infections and about 10-fold for high MOI infections (Fig 1B). All of these differences were statistically significant. Taken together, this data suggest that depletion of dynamin causes severe growth impairment of MCMV.

**Figure 1.**
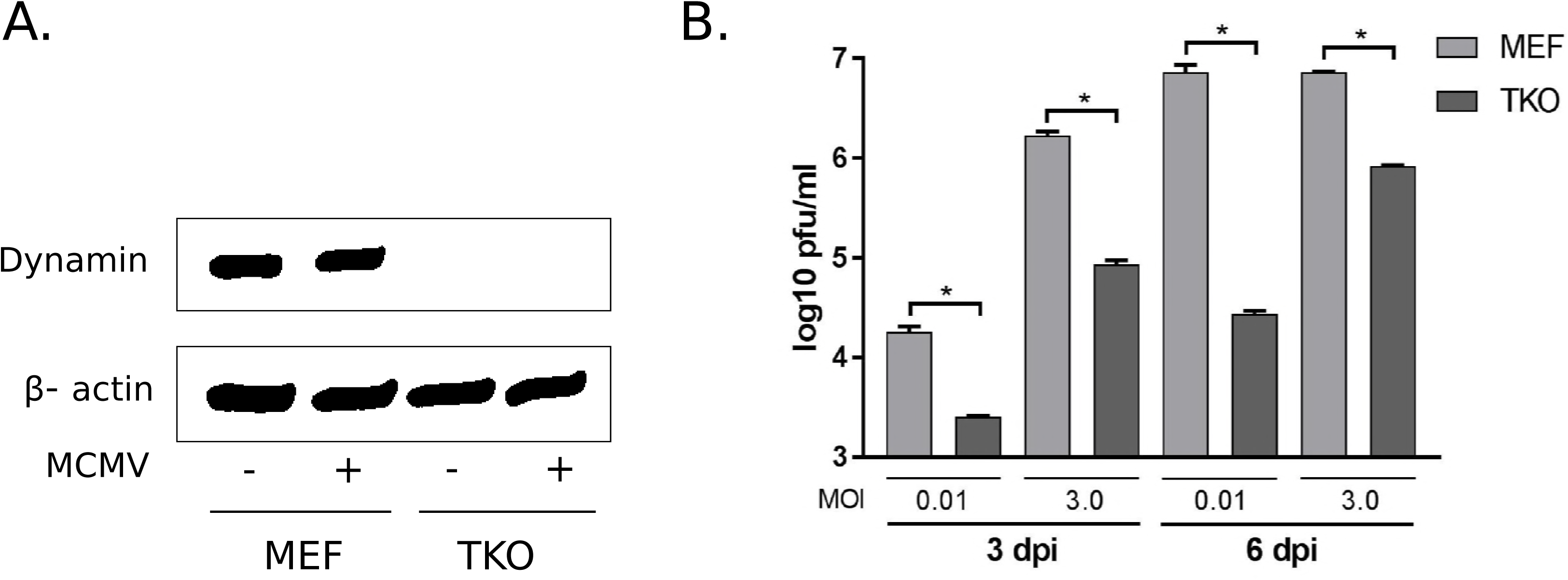
Dynamin depletion impacts the growth of MCMV in fibroblasts. Dynamin was depleted in engineered mouse embryonic fibroblasts (MEF) by a tamoxifen (4-hydroxytamoxifen; 4-HT) inducible knockout strategy leading to the generation of triple dynamin knockout (TKO) cells as described earlier (32). MEFs were treated with 4-HT for 2 days and then 4-HT containing media was replaced with fresh 4-HT-free media and cells were incubated for additional 2 days. A) Parental MEF and TKO cells were infected with MCMV K181 strain at MOI 3.0 (+) or mock-infected (-) and cell lysates were harvested at 4 hours post infection for immunoblot probing for dynamin. β-actin was used as a loading control. B) Parental MEF and TKO cells were infected with MCMV-K181 strain at MOI 3.0 or 0.01 and cells with media were harvested at three- or six-days post infection before plating for virus titers on wild type MEFs. Triplicate samples were used in experiments. A two-tailed unpaired t-test with Welch’s correction (unequal variance assumption) was used for statistical analysis of differences. P Values <0.05 were considered significant (*). dpi: days post infection.

### Dynamin depletion interferes with MCMV entry in fibroblasts and affects late protein expression

To explore the impact of dynamin on virus entry, parental MEF and TKO cells were infected with MCMV-K181 at MOI 3 and cell lysates were analyzed for expression of immediate early (IE1; m123) protein. At 4- and 24 hours post infection (hpi), IE1 was detected in both MEF and TKO; however, levels of IE1 were significantly reduced in TKO (Fig 2) indicating that virus entry was affected but not completely eliminated in TKO cells. Similarly, reduced levels of early protein m06 were detected in TKO compared to MEF at 24 hpi. Expression levels were detected to be similar in TKO and MEF for both IE1 and m06 at 48 hpi. In contrast, late viral protein m55 was expressed at lower levels in TKO cells at 48 hpi. Altogether, these data indicate that MCMV enters less efficiently in dynamin-depleted fibroblasts but establishes infection, albeit with a compromised expression of late viral proteins.

**Figure 2.**
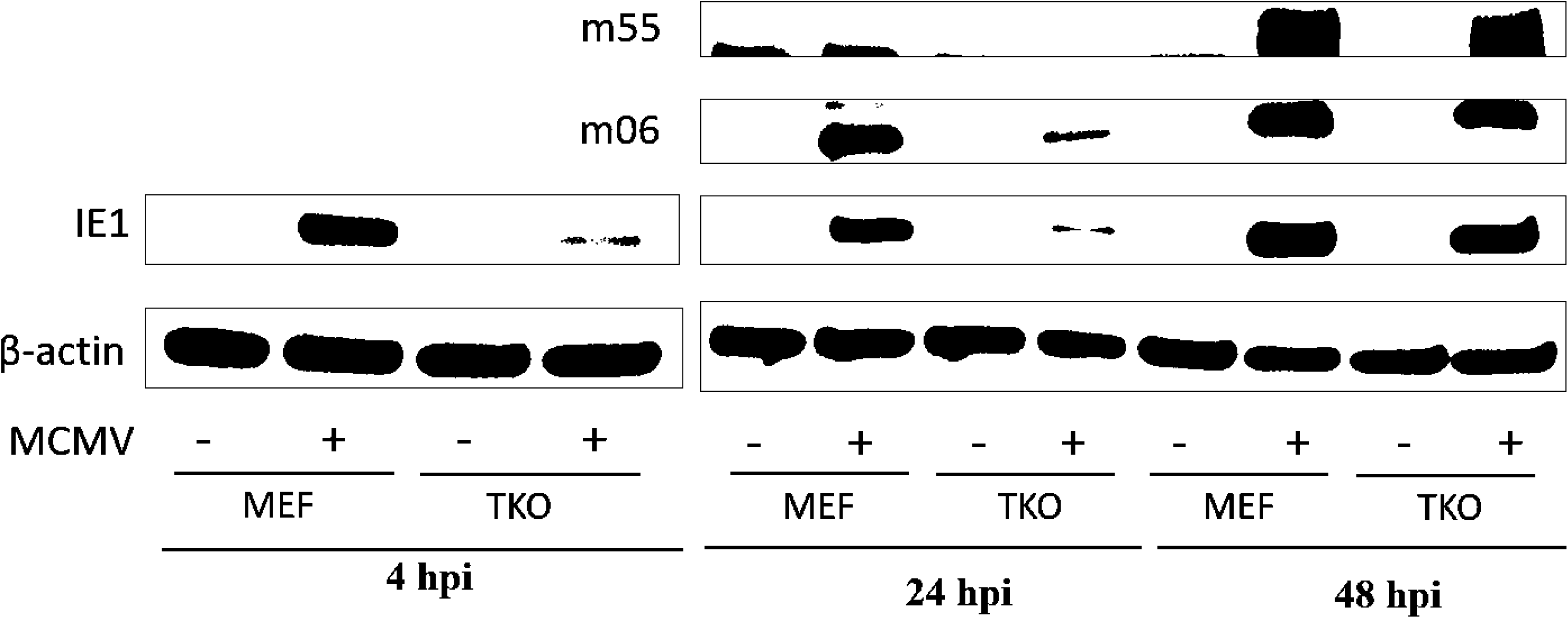
Dynamin depletion reduces the entry of MCMV in fibroblasts and interferes with late protein expression. Parental MEFs and TKO cells were infected with MCMV-K181 at MOI 3.0 (+) or mock-infected (-). Cells were harvested at 4 h, 24h and 48 h post infection and probed for immediate early (IE1; m123), early (m06), and late (m55) viral proteins. β-actin served as loading control. hpi: hours post infection.

There is a certain possibility that inefficient virus entry in TKO cells, as evidenced by reduced IE1 expression at early times post infection (Fig. 2), leads to an overall delayed replication cycle. To probe this further, we performed a full single-step virus growth curve analysis. Parental MEF and TKO cells were either mock-infected or infected with MCMV-K181 at MOI 3.0 and cells and medium were harvested followed by quantification of plaque forming units. As summarized in Figure 3A, viral growth defect in TKO cells was evident as early as 3 days post infection (dpi) and continued up to 7 dpi. We performed a cell viability test of mock and MCMV-K181 infected MEF and TKO up to 7 dpi in parallel to rule out the possibility that this growth defect could be due to a viability disadvantage in TKO cells (Fig 3B). Uninfected MEF and TKO cells were >95% viable until 5 days in cell culture. At 7 days, TKO cells showed more cell death compared to MEF. During MCMV infection, both TKO and MEF cells showed significant cell death starting at 1 dpi; however, TKO cells showed more resistance to MCMV induced cell-death, especially at 3- and 5-days post infection.

**Figure 3.**
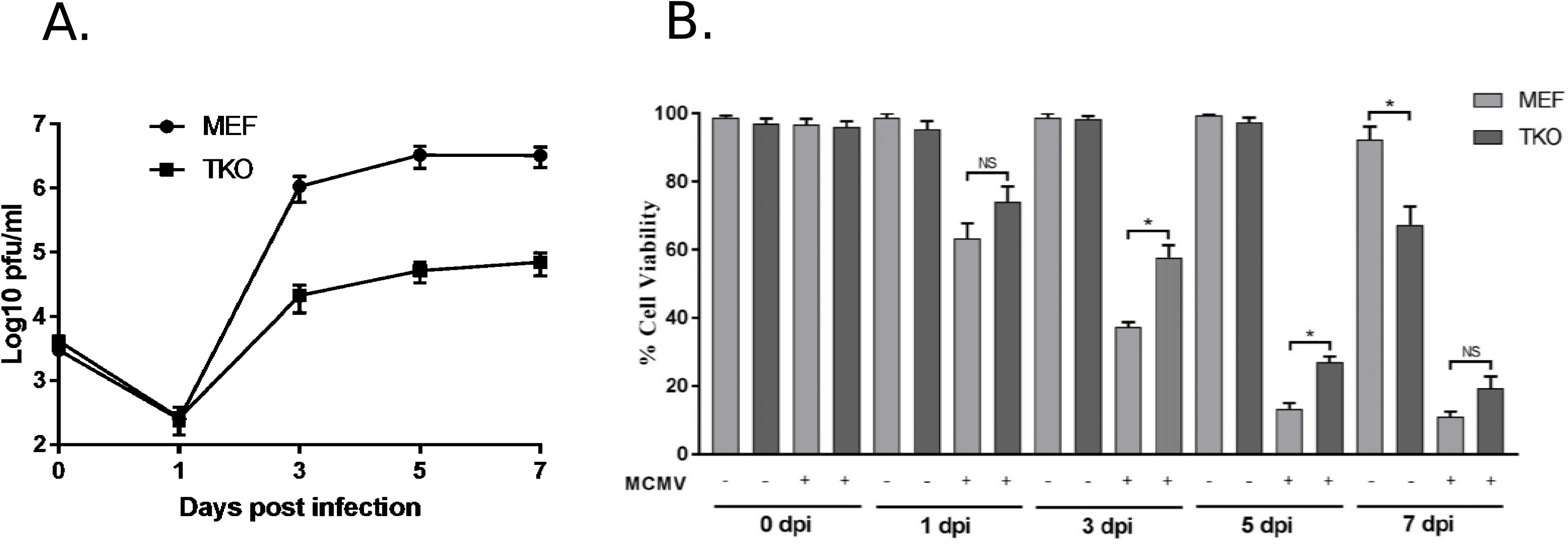
Impact of dynamin depletion on MCMV growth and cell viability. A) Parental MEF and TKO cells were infected with MCMV-K181 at MOI 3.0 and cells with media were harvested at zero to 7 days post infection followed by estimation of virus titers on wild type MEFs. B) Parental MEF and TKO cells were infected with MCMV-K181 at MOI 3.0 (+) or mock-infected (-) and cell viability at the indicated time points was determined by trypan blue exclusion assay. Triplicate samples were used in experiments. A two-tailed unpaired t-test with Welch’s correction (unequal variance assumption) was used for statistical analysis of differences. P Values <0.05 were considered significant (*).

To probe whether the observed growth defect in TKO cells (Fig 3A) reflects a defect in virus replication or release from cells, we separated cells and supernatants at different times post infection and evaluated the viral titers (Fig. 4). The results indicate significant reduction in both cell and supernatant-associated virus in TKO cells at 3 dpi for both low and high MOI. A similar trend was observed at 6 dpi for low MOI. Interestingly, at high MOI, the supernatant titers for TKO cells were significantly reduced but cell-associated titers were equivalent to MEF cells at 6 dpi. In summary, the data corroborate the results from growth analysis (Fig 1, 3) and protein expression studies (Fig 2) that virus growth is delayed in TKO cells; however, it also indicates that virus growth in TKO cells catches up with WT-MEF at late time post infection and the growth defect observed at this time is almost entirely due to a defect in virus release.

**Figure 4.**
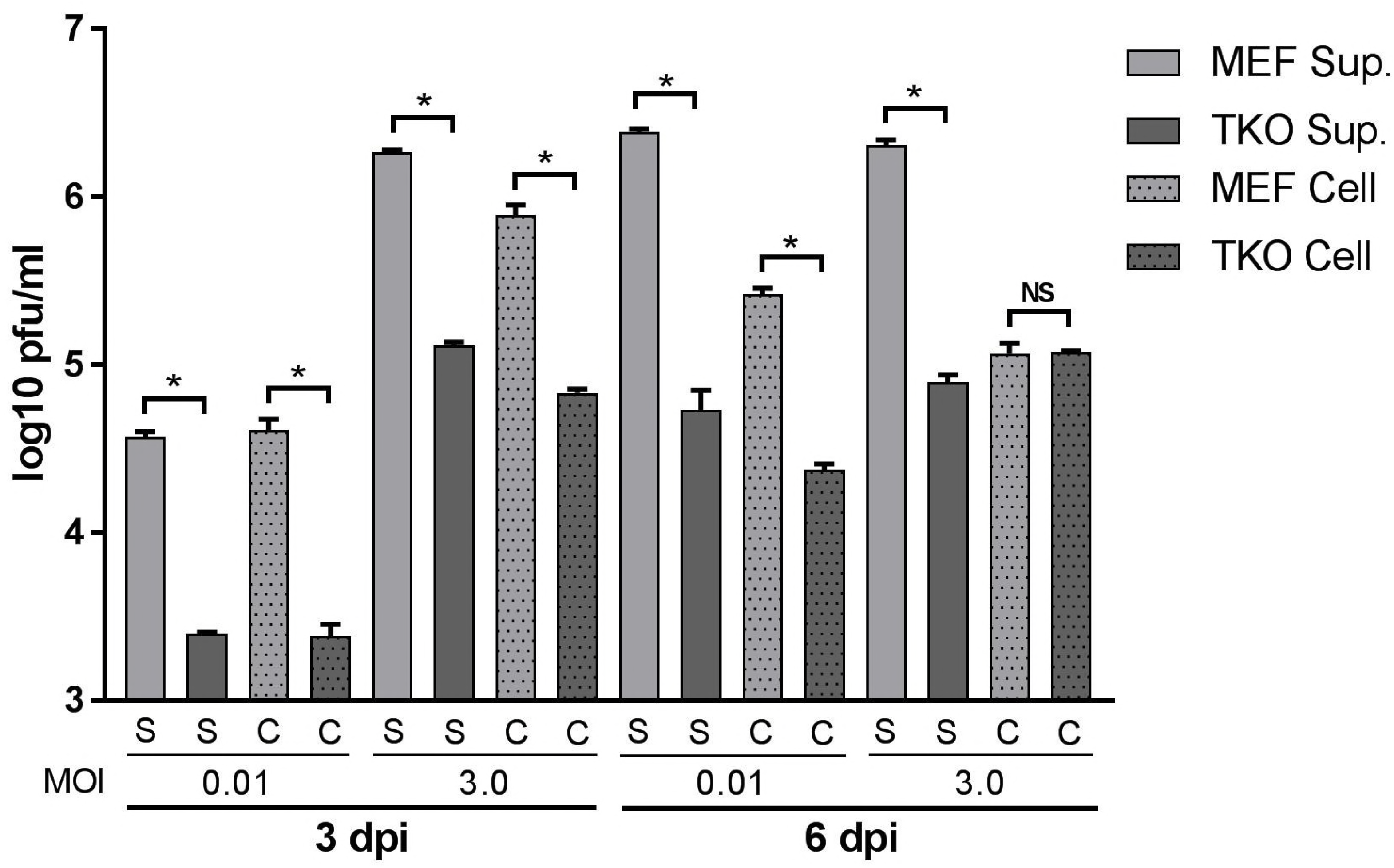
Dynamin depletion reduces both cell-associated cell-free virus levels. Parental MEF and TKO cells were infected with MCMV-K181 at MOI 3.0 or 0.01 and cells with media were harvested at three- or six-days post infection. Media and cells were separated by low-speed centrifugation and viral loads in supernatant (S) and cells (C) were quantified by titering on wild-type MEF. Triplicate samples were used in experiments. A two-tailed unpaired t-test with Welch’s correction (unequal variance assumption) was used for statistical analysis of differences. P Values <0.05 were considered significant (*).

### Lack of dynamin does not impair the localization of early and late viral proteins

In order to understand the impact of dynamin depletion on virus protein trafficking, we examined the localization of MCMV early and late proteins in MEF and TKO cells at 48 hpi by immunofluorescence assay (IFA). MCMV immediate early (IE1; m123) protein was expressed and localized to the nucleus in both parental MEF and TKO cells (Fig 5, top 2 panels). MCMV early (m04) protein was expressed in the cytoplasm but concentrated around the nuclear periphery in both cell types (Fig 5, middle 2 panels). Similarly, MCMV late (m55) protein was expressed in the cytoplasm of both cells in diffuse as well as punctate (possibly virion associated) forms (Fig 5, bottom 2 panels). Collectively, these data indicate that dynamin-depletion does not affect the expression and localization of early to late viral proteins that is evident in IFA.

**Figure 5.**
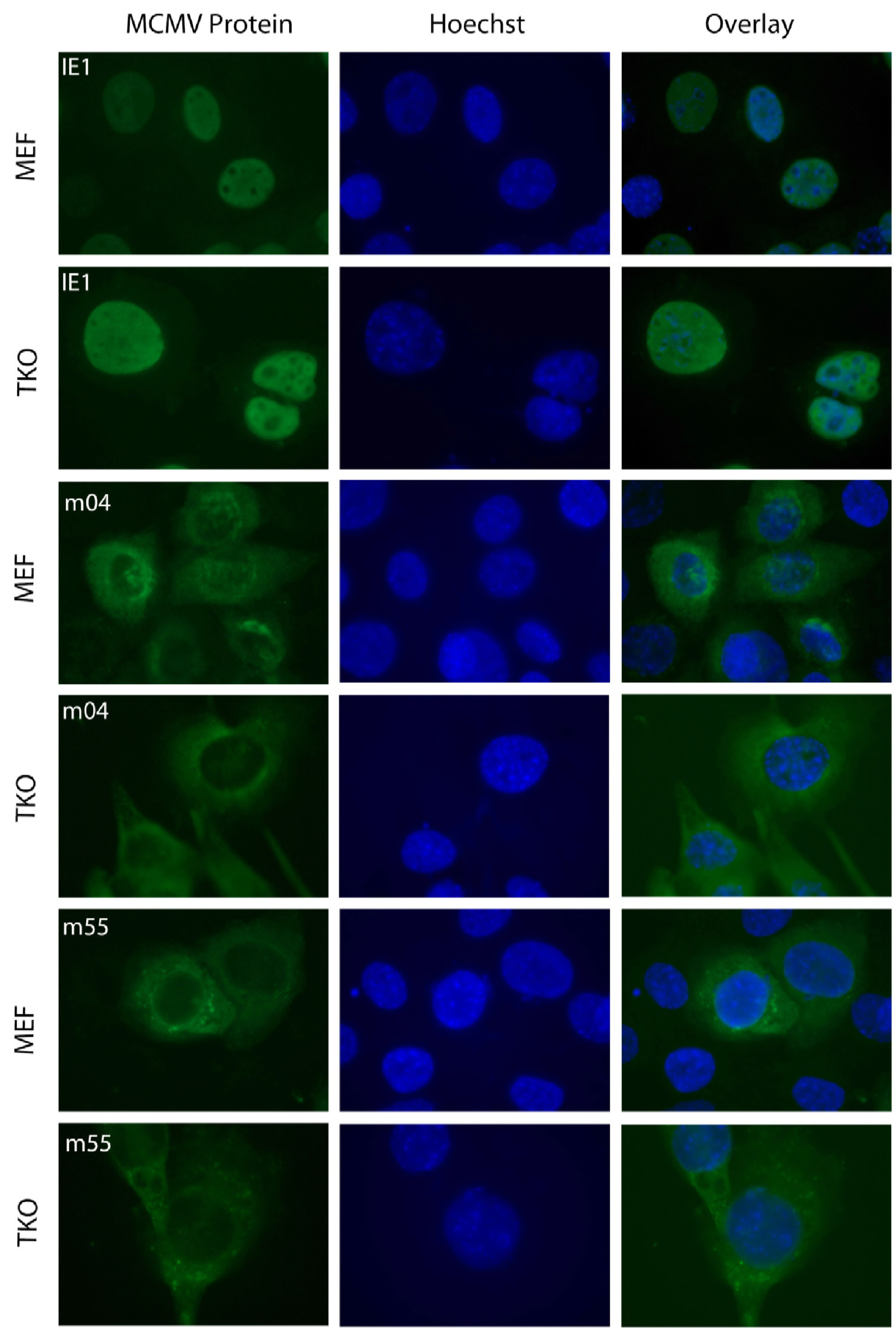
Dynamin depletion does not affect the localization of early and late viral proteins in infected cells. Parental MEF and TKO cells were infected with MCMV-K181 at MOI 3.0, fixed for IFA at 48 hours post infection and stained for MCMV immediate early (IE1; m123), early (m04), and late (m55) proteins. MCMV proteins labeled as green (left column), DNA (in nuclei) detected by Hoechst (middle column) and composite images (overlay, right column) from the same field of each panel are shown. IE1 protein localized to the nuclear compartment whereas m04 and m55 proteins localized to the cytoplasm concentrating at the periphery of the nucleus in both MEF and TKO cells.

### Lack of dynamin affects the formation of vAC

vAC is known to be the site of cytoplasmic virus maturation. Since the growth data (Fig 4) showed a defect in virus release at late times post infection, we investigated the formation of vAC in TKO and MEF cells to analyze any defects that would translate to a defect in virus envelopment and release. Mock-infected MEF showed the presence of perinuclear Golgin-97 staining, consistent with the presence of Golgi-stacks (36) (Fig 6). Similar perinuclear Golgin-97 staining was also observed in mock-infected TKO cells. Infected MEF showed a perinuclear Goglin-97 ring formation, as observed in HCMV infected fibroblasts and marks the vAC (12, 37). In infected TKO cells, the Golgin-97 accumulated in the perinuclear region but none of the cells examined (>100) showed the typical ring formation. Thus, the data indicate that assembly of vAC is compromised in TKO cells.

**Figure 6.**
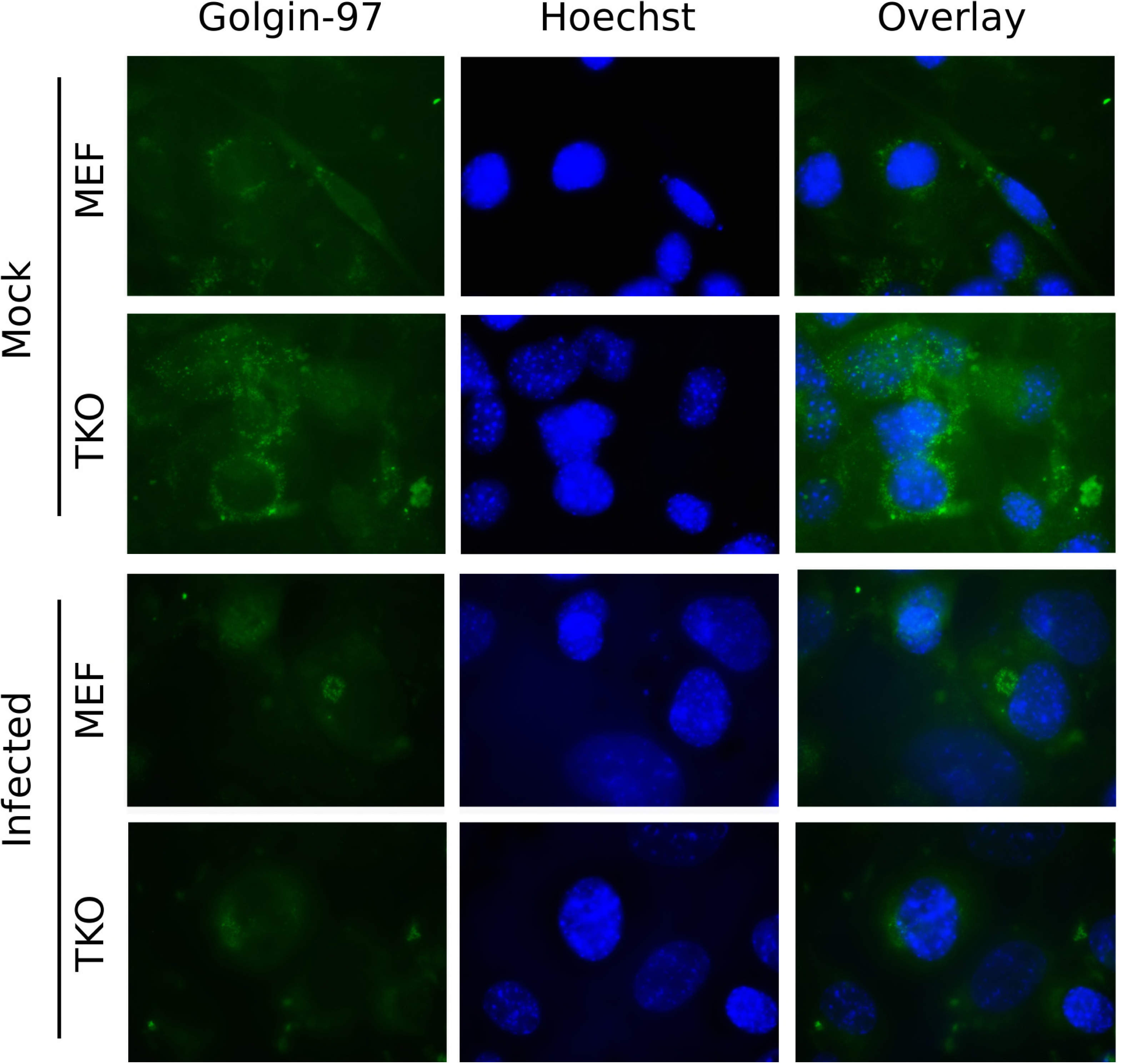
Formation of vAC is compromised in dynamin depleted cells. Parental MEF and TKO cells were mock-infected or infected with MCMV-K181 at MOI 3.0, fixed for IFA at 48 hours post infection and stained for Golgin-97. Golgin-97 labeled as green (left column), DNA (in nuclei) detected by Hoechst (middle column) and composite images (overlay, right column) from the same field of each panel are shown.

### MCMV nuclear stages are intact in dynamin-depleted fibroblasts but cytoplasmic virus maturation is significantly impaired

Parental MEF and TKO cells were infected with MCMV-K181 at an MOI of 3.0. At 72 hours post infection, cells were fixed for processing and imaging under transmission electron microscope. Both cell types showed typical infected cells morphology with a kidney-bean shaped nucleus and the presence of nuclear and cytoplasmic inclusions (Fig 7A, E). The nucleus of both MEF and TKO cells contained all three types of capsids (A (empty), B (scaffold-containing) and C (DNA-containing)) reported for herpesviruses (Fig 7B, F) (8). Quantification of these capsid types revealed similar proportions in both cell types (Fig 7I) indicating intact nucleocapsid maturation. In contrast, very few virus particles were observed in the cytoplasm of TKO cells (Fig 7G, H); however, these particles contained genomic DNA and presence of tegument proteins could be appreciated on the surface of these capsids (Fig 7G inset). Virus envelopment was not evident in TKO cells. Several enveloping (Fig 7C inset) or enveloped virus particles were present in the cytoplasm of parental MEF (Fig 7C, D). Another striking difference was the presence of intact Golgi stacks in TKO cells (Fig 7 G, H), which were fragmented to different degrees in MEF cells (Fig 7 C, D) indicating increased vesiculation. Examination of cytoplasm revealed no virions or partially enveloped particles in TKO cells (Fig 7 G, H, J) in contrast to significant number of virions and enveloping particles in the parental MEF (Fig. 7 C, D, J). In summary, the data indicate that cytoplasmic maturation is severely compromised in TKO cells.

**Figure 7.**
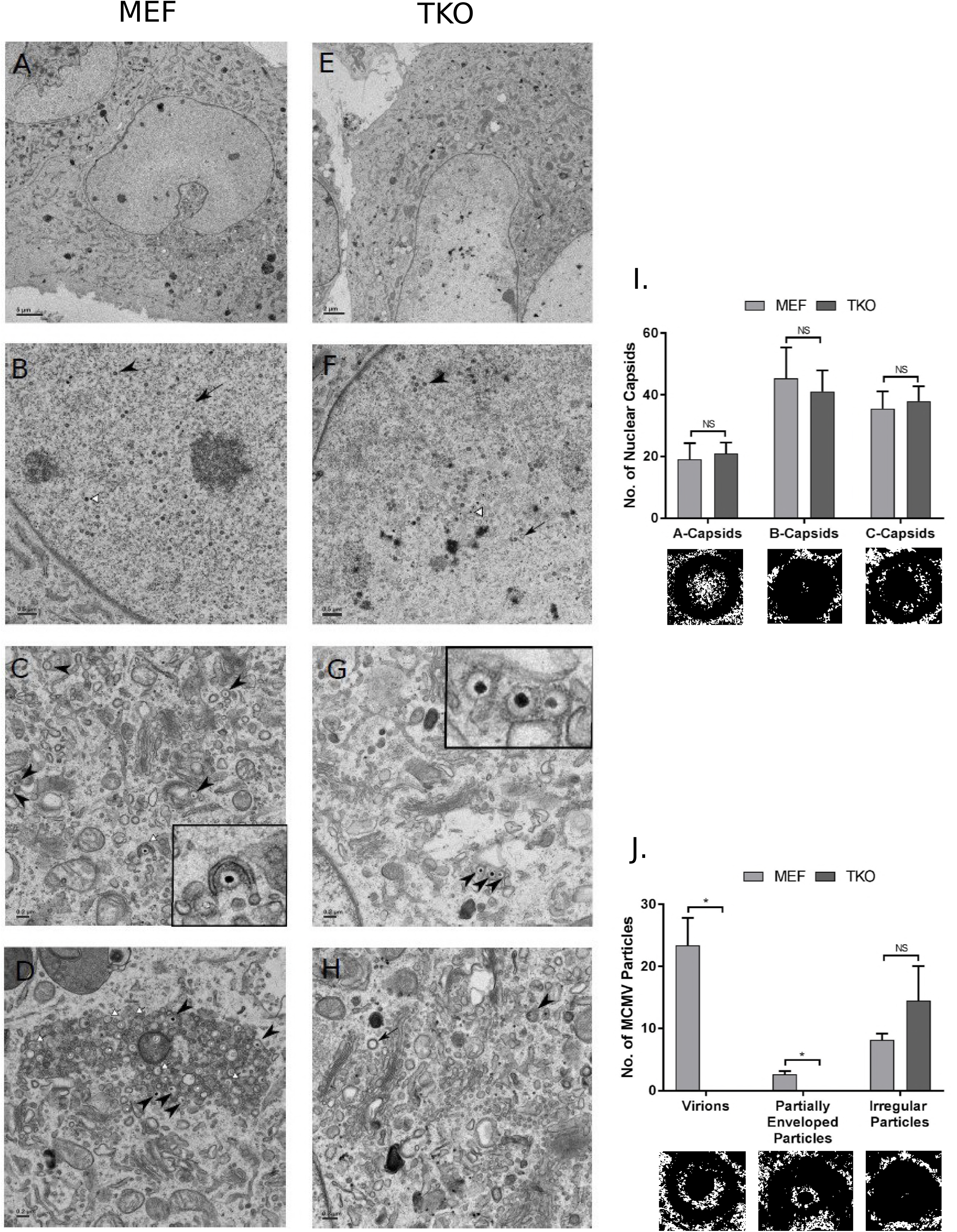
MCMV nuclear stages are intact in TKO cells but cytoplasmic virus maturation is significantly impaired. Transmission electron micrographs (TEM) of MEF (A-D) and TKO (E-H) cells infected with MCMV-K181. Cells were infected at an MOI of 3.0 and fixed for processing at 3 days post infection. (A and E) A single infected cell showing nucleus as well as cytoplasm. (B and F) Infected cell nucleus illustrating A-(black arrows), B-(black arrowheads), and C-(white triangular arrows) capsids. (C, D, G and H) Cytoplasmic section illustrating DNA containing capsids (black arrowheads), partially enveloped/enveloping capsids (white arrows) and virus-like particles that are difficult to type morphologically (black arrow). The inset in C magnifies and illustrates a partially enveloped virion and the inset in G illustrates non-enveloped DNA containing capsids that appear to have some intact tegument. Intact Golgi stacks can be appreciated in G and H, whereas fragmented Golgi is prominent in C and D. I) Nuclear capsids were quantified in MEF (n=6) and TKO (n=7) cells. Representative of each type of capsid is shown under the graph. J) MCMV particles in the cytoplasm of MEF (n=8) and TKO (n=7) cells were quantified. A representative example of virion, partially enveloped particle and irregular particle is shown under the graph. A two-tailed unpaired t-test with Welch’s correction (unequal variance assumption) was used for statistical analysis of differences. P Values <0.05 were considered significant (*).

### Dynasore-treated cells mimic the dynamin knockout phenotype

To rule out any unknown peculiarity in TKO cells that may be responsible for the virus maturation defects evident in these cells, we treated wild-type primary MEF with 50 µM of an established dynamin inhibitor (dynasore) and subsequently infected with MCMV-K181 to study virus entry and growth. Dynasore is a small molecule that is well established to specifically abolish dynamin activity in cells without an impact on cell viability (31). Dynasore-treatment resulted in a decrease in IE1 gene expression at 4 hours post infection but this expression was normalized at 48 hours post infection (Fig 8A) similar to the results obtained for TKO cells (Fig 2). Analysis of virus growth at low (0.05) and high (3.0) MOI indicated significant differences between dynasore-treated and mock-treated cells (Fig 8B). These results also correlate with the results obtained for TKO cells (Fig 1, 3). Thus, the phenotype we observed in TKO cells is indeed due to the deficiency of dynamin function and is not an aberrant effect of dynamin depletion on a single cell type.

**Figure 8.**
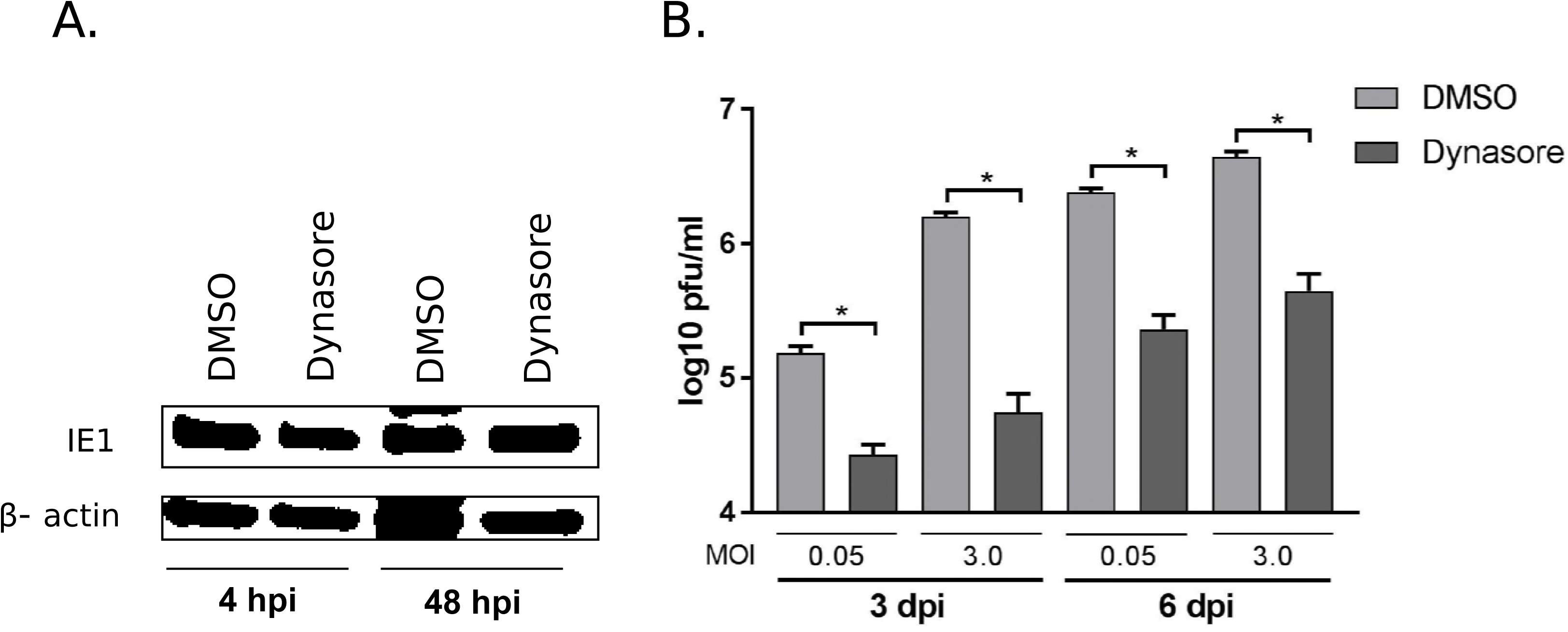
Dynasore-treated MEFs allow virus entry and gene expression but compromise virus growth. A) MEF were treated with dynasore (50 µM) or mock (DMSO) and infected with MCMV-K181 at an MOI of 3.0. Cell lysates were harvested at 4 h and 48 h post infection for immunoblot probing for MCMV IE1 protein. β-actin was used as a loading control. B) Dynasore or mock treated cells were infected with MCMV-K181 at MOI 0.05 or 3.0 and cells were harvested at three- or six-days post infection before plating for virus titers on wild type MEFs. Triplicate samples were used in experiments. A two-tailed unpaired t-test with Welch’s correction (unequal variance assumption) was used for statistical analysis of differences. P Values <0.05 were considered significant. hpi: hours post infection, dpi: days post infection.

## Discussion

Endosomal membranes have been implicated in herpesvirus maturation; however, the role of specific endocytic pathways in herpesvirus morphogenesis remains largely unexplored. In the current work, we show that dynamin-mediated endocytic pathways are important for CMV maturation. We utilized recently characterized triple dynamin knockout cells for these studies to provide convincing evidence that these pathways are important at a late stage of CMV life cycle that involves virus morphogenesis, gain of infectivity and egress of mature particles.

The current studies were influenced by our earlier studies on HCMV where we utilized laboratory strains that utilize a glycoprotein-mediated fusion mechanism at plasma membrane to enter the cells instead of endocytosis (30). The HCMV entry pathways in different cells types have been studied in detail and it is well known that laboratory strains enter the cells via a pH-independent fusion mechanism at the plasma membrane (38, 39). We used clathrin and dynamin inhibitors in the above study to reveal a role of endocytic processes on HCMV maturation. The data from these studies indicated an impact of pharmacological inhibition of dynamin-clathrin pathways on HCMV maturation; however, virus entry and early gene expression remained intact. To be able to extend the study of virus biology in an appropriate animal model, we utilized an established dynamin-knockout mouse cell model that has been extensively characterized (32, 40, 41) and is free from any side-effects that chemical inhibitors may have on cells. It also provides the ability to test MCMV instead of HCMV, which would be useful for future *in vivo* studies looking to characterize the effect of endocytic inhibitors in a mouse model of CMV infection. This is important because dynamin inhibitors have already shown a therapeutic potential against several infectious agents (reviewed in (33)).

After successfully establishing a near-complete depletion of dynamin in TKO cells, we measured its impact on MCMV growth and yield. Dynamin depletion had significant impact on virus growth at low as well as high multiplicity of infection. Although analysis of early viral gene expression revealed an impact on early time of infection (4 h and 24h), these differences were normalized by 48 h indicating that the reduction in virus growth in TKO cells observed at late times post infection is unlikely due to defects in virus entry or early gene expression. To further investigate this point, we analyzed the expression and distribution of early to late viral proteins in infected cells. Expression of early proteins (IE1, m06) was at equivalent levels at 48 hpi; however, the late protein (m55) expression was significantly reduced indicating that the defects in CMV replication in dynamin depleted cells relate to late steps in virus replication that include the expression of late genes. Localization of viral proteins (IE1, m04 and m55) in TKO cells was not significantly different from paternal MEF. These results rule out the possibility that virus growth defects could be due to an impact on early viral gene expression or abnormal localization of major viral proteins in dynamin depleted cells. Investigation of cellular protein (Golgin97) localization revealed that vAC does not form properly in TKO cells; therefore, cytoplasmic virus maturation and egress are likely impaired.

The possibility of a defect at a late stage of virus maturation was analyzed by ultrastructural detailed analysis of infected cells. Nuclear stages of virus replication including capsid assembly and DNA packaging were intact based on the numbers and types of capsid particles present in the nuclei of TKO versus parental MEF. Equivalent proportions of A, B and C capsid forms (8) were observed in both cell types (Fig 7I). Most importantly, the presence of similar numbers of C-capsids (DNA packaged capsids) in two cell types indicate that dynamin did not influence the stages of virus replication up to the point of production of DNA packaged capsids, which go on to become infectious virions. Thus, the defects would either be at the nuclear egress of packaged capsids or at the cytoplasmic stage of virus maturation and egress. Further, a block at nuclear egress was ruled out on the basis of the absence of any large buildup of assembled particles at the inner nuclear membrane in ultrastructural images (Fig 7E, F). Moreover, a few virus particles were present in the cytoplasm of TKO cells indicating that nuclear egress could not be completely blocked. These cytoplasmic virus particles in TKO cells were mostly unenveloped (Fig 7G) or appeared to be morphologically abnormal/degrading (Fig 7H, arrowhead). This is not unusual since an absence of an envelope would ultimately lead to degradation of naked virus particles in the cytoplasm. We saw a similar phenotype of capsids degrading in the cytoplasm of pp150 mutant virus infected cells in our earlier study (37). This degradation happened despite the re-localization of viral DNA from the nucleus to the cytoplasm (42). Since viral DNA cannot exit the nucleus independent of virus capsids (and even if it did, it would degrade rapidly due to strong cytoplasmic nucleases), the most convincing explanation is that capsids carry viral DNA to the cytoplasm but are unable to maintain their integrity in the absence of essential inner tegument proteins.

Exocytic vesicles and vesicles containing clathrin-coated pits are generated from Golgi fragmentation. Anti-dynamin antibodies have been shown to block the formation of these vesicles in a cell-free assay (43). This vesiculation is restored upon addition of purified dynamin. Thus, it comes as no surprise that dynamin depleted cells have more intact Golgi stacks compared to parental cells when infected by CMV. This vesiculation may contribute significantly to CMV envelopment, which is compromised in dynamin-depleted cells. The late virus maturation defect observed in the current study, along with an evident disruption of vAC formation point towards the important role of dynamin-mediated vesicular pathways in CMV maturation (Fig 9).

**Figure 9.**
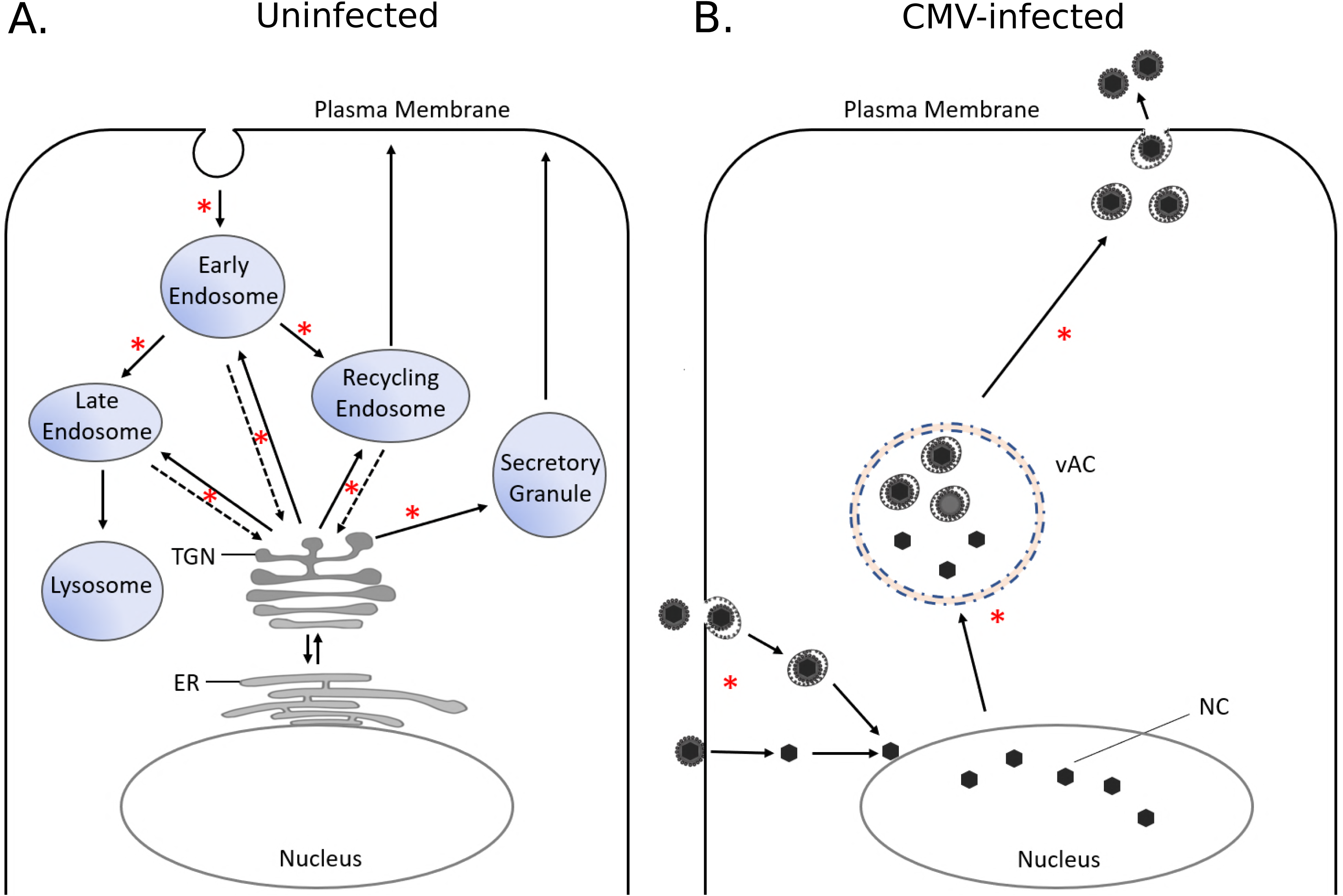
Proposed model for the functions of dynamin in CMV-infected cells. A) Dynamin plays a role in membrane remodeling at different stages of endosomal trafficking. Newly synthesized proteins in the ER are sorted in the TGN targeted for their final destination in the cell or secreted forms. TGN also receives input from the endocytic pathways (broken arrows) where dynamin is implicated. B) Proposed model of function of dynamin in a CMV-infected cell. Proposed point of critical activity of dynamin is marked with an asterisk. ER: endoplasmic reticulum, TGN: trans-Golgi network, NC: Nuclear capsid, vAC: Virus assembly complex.

There is little doubt that endosomal systems contribute to the process of herpesvirus maturation; however, examples of specific virus proteins hijacking host endocytic machinery are lacking. A study based on mass spectroscopic analysis of protein interactions in HCMV-infected cells indicated that the tegument protein pp150 directly interacts with clathrin (44). pp150 has established roles in virus maturation (37) and it is certainly possible that this pp150-clathrin interaction is functional during virus maturation and egress. More specific interactions of herpesvirus proteins with endocytic systems are likely to be revealed by studying host factors such as dynamin that are important in the late stages of herpesvirus maturation.

## Acknowledgments

We are thankful to Pietro De Camilli at Yale University for the gift of triple dynamin-knockout (TKO) cells. Hong Yi at the Robert P. Apkarian Integrated Electron Microscopy Core at Emory University acquired the electron microscopy data. The research was supported by American Heart Association Scientist Development Grant (Award 14SDG20390009, PI: Tandon).

## Author Contributions

RT designed the experiments; MHH, LED, RPM and RT performed the experiments and analyzed the data; RKB and DM helped with virus growth assays and plaque counting. RT wrote and edited the manuscript.

## References

1. Mercer J, Schelhaas M, Helenius A. 2010. Virus entry by endocytosis. Annu Rev Biochem 79:803–833.

2. Schelhaas M. 2010. Come in and take your coat off - how host cells provide endocytosis for virus entry. Cell Microbiol 12:1378–1388.

3. Sun Y, Tien P. 2013. From endocytosis to membrane fusion: emerging roles of dynamin in virus entry. Crit Rev Microbiol 39:166–179.

4. Humphries AC, Way M. 2013. The non-canonical roles of clathrin and actin in pathogen internalization, egress and spread. Nat Rev Microbiol 11:551–560.

5. Blanchard E, Belouzard S, Goueslain L, Wakita T, Dubuisson J, Wychowski C, Rouille Y. 2006. Hepatitis C virus entry depends on clathrin-mediated endocytosis. J Virol 80:6964–6972.

6. Mocarski ES, Jr., Shenk, T., Pass R. F. 2006. Cytomegaloviruses., p. 2701–2772. In D M Knipe and P M Howley (ed), Fields Virology 5th Edition Lippincott Williams & Wilkins, Philadelphia.

7. Hellberg T, Passvogel L, Schulz KS, Klupp BG, Mettenleiter TC. 2016. Nuclear Egress of Herpesviruses: The Prototypic Vesicular Nucleocytoplasmic Transport. Adv Virus Res 94:81–140.

8. Tandon R, Mocarski ES. 2012. Viral and host control of cytomegalovirus maturation. Trends Microbiol 20:392–401.

9. Henaff D, Radtke K, Lippe R. 2012. Herpesviruses exploit several host compartments for envelopment. Traffic 13:1443–1449.

10. Buckingham EM, Jarosinski KW, Jackson W, Carpenter JE, Grose C. 2016. Exocytosis of Varicella-Zoster Virus Virions Involves a Convergence of Endosomal and Autophagy Pathways. J Virol 90:8673–8685.

11. Owen DJ, Crump CM, Graham SC. 2015. Tegument Assembly and Secondary Envelopment of Alphaherpesviruses. Viruses 7:5084–5114.

12. Das S, Vasanji A, Pellett PE. 2007. Three-dimensional structure of the human cytomegalovirus cytoplasmic virion assembly complex includes a reoriented secretory apparatus. J Virol 81:11861–11869.

13. Das S, Pellett PE. 2011. Spatial relationships between markers for secretory and endosomal machinery in human cytomegalovirus-infected cells versus those in uninfected cells. J Virol 85:5864–5879.

14. Karleusa L, Mahmutefendic H, Tomas MI, Zagorac GB, Lucin P. 2017. Landmarks of endosomal remodeling in the early phase of cytomegalovirus infection. Virology 515:108–122.

15. Tandon R, AuCoin DP, Mocarski ES. 2009. Human cytomegalovirus exploits ESCRT machinery in the process of virion maturation. J Virol 83:10797–10807.

16. Chiu YF, Sugden B, Chang PJ, Chen LW, Lin YJ, Lan YC, Lai CH, Liou JY, Liu ST, Hung CH. 2012. Characterization and intracellular trafficking of Epstein-Barr virus BBLF1, a protein involved in virion maturation. J Virol 86:9647–9655.

17. Crump CM, Yates C, Minson T. 2007. Herpes simplex virus type 1 cytoplasmic envelopment requires functional Vps4. J Virol 81:7380–7387.

18. Brunetti CR, Dingwell KS, Wale C, Graham FL, Johnson DC. 1998. Herpes simplex virus gD and virions accumulate in endosomes by mannose 6-phosphate-dependent and -independent mechanisms. J Virol 72:3330–3339.

19. Tooze J, Hollinshead M, Reis B, Radsak K, Kern H. 1993. Progeny vaccinia and human cytomegalovirus particles utilize early endosomal cisternae for their envelopes. Eur J Cell Biol 60:163–178.

20. Beitia Ortiz de Zarate I, Kaelin K, Rozenberg F. 2004. Effects of mutations in the cytoplasmic domain of herpes simplex virus type 1 glycoprotein B on intracellular transport and infectivity. J Virol 78:1540–1551.

21. Van Minnebruggen G, Favoreel HW, Nauwynck HJ. 2004. Internalization of pseudorabies virus glycoprotein B is mediated by an interaction between the YQRL motif in its cytoplasmic domain and the clathrin-associated AP-2 adaptor complex. J Virol 78:8852–8859.

22. Tugizov S, Maidji E, Xiao J, Pereira L. 1999. An acidic cluster in the cytosolic domain of human cytomegalovirus glycoprotein B is a signal for endocytosis from the plasma membrane. J Virol 73:8677–8688.

23. Hollinshead M, Johns HL, Sayers CL, Gonzalez-Lopez C, Smith GL, Elliott G. 2012. Endocytic tubules regulated by Rab GTPases 5 and 11 are used for envelopment of herpes simplex virus. EMBO J 31:4204–4220.

24. Schauflinger M, Fischer D, Schreiber A, Chevillotte M, Walther P, Mertens T, von Einem J. 2011. The tegument protein UL71 of human cytomegalovirus is involved in late envelopment and affects multivesicular bodies. J Virol 85:3821–3832.

25. Praefcke GJ, McMahon HT. 2004. The dynamin superfamily: universal membrane tubulation and fission molecules? Nat Rev Mol Cell Biol 5:133–147.

26. Pigino G, Morfini GA, Brady TS. 2012. Intracellular Trafficking. Basic Neurochemistry (Eighth Edition):119–145.

27. Anggono V, Robinson PJ. 2009. Dynamin. Encyclopedia of Neuroscience:725–735.

28. Kirchhausen T. 1998. Vesicle formation: dynamic dynamin lives up to its name. Curr Biol 8:R792–794.

29. Mettlen M, Pucadyil T, Ramachandran R, Schmid SL. 2009. Dissecting dynamin’s role in clathrin-mediated endocytosis. Biochem Soc Trans 37:1022–1026.

30. Archer MA, Brechtel TM, Davis LE, Parmar RC, Hasan MH, Tandon R. 2017. Inhibition of endocytic pathways impacts cytomegalovirus maturation. Sci Rep 7:46069.

31. Macia E, Ehrlich M, Massol R, Boucrot E, Brunner C, Kirchhausen T. 2006. Dynasore, a cell-permeable inhibitor of dynamin. Dev Cell 10:839–850.

32. Park RJ, Shen H, Liu L, Liu X, Ferguson SM, De Camilli P. 2013. Dynamin triple knockout cells reveal off target effects of commonly used dynamin inhibitors. J Cell Sci 126:5305–5312.

33. Harper CB, Popoff MR, McCluskey A, Robinson PJ, Meunier FA. 2013. Targeting membrane trafficking in infection prophylaxis: dynamin inhibitors. Trends Cell Biol 23:90–101.

34. Badea TC, Wang Y, Nathans J. 2003. A noninvasive genetic/pharmacologic strategy for visualizing cell morphology and clonal relationships in the mouse. J Neurosci 23:2314–2322.

35. Zurbach KA, Moghbeli T, Snyder CM. 2014. Resolving the titer of murine cytomegalovirus by plaque assay using the M2-10B4 cell line and a low viscosity overlay. Virol J 11:71.

36. Bardin S, Miserey-Lenkei S, Hurbain I, Garcia-Castillo D, Raposo G, Goud B. 2015. Phenotypic characterisation of RAB6A knockout mouse embryonic fibroblasts. Biol Cell 107:427–439.

37. Tandon R, Mocarski ES. 2008. Control of cytoplasmic maturation events by cytomegalovirus tegument protein pp150. J Virol 82:9433–9444.

38. Vanarsdall AL, Johnson DC. 2012. Human cytomegalovirus entry into cells. Curr Opin Virol 2:37–42.

39. Ryckman BJ, Jarvis MA, Drummond DD, Nelson JA, Johnson DC. 2006. Human cytomegalovirus entry into epithelial and endothelial cells depends on genes UL128 to UL150 and occurs by endocytosis and low-pH fusion. J Virol 80:710–722.

40. Shen H, Ferguson SM, Dephoure N, Park R, Yang Y, Volpicelli-Daley L, Gygi S, Schlessinger J, De Camilli P. 2011. Constitutive activated Cdc42-associated kinase (Ack) phosphorylation at arrested endocytic clathrin-coated pits of cells that lack dynamin. Mol Biol Cell 22:493–502.

41. Antonny B, Burd C, De Camilli P, Chen E, Daumke O, Faelber K, Ford M, Frolov VA, Frost A, Hinshaw JE, Kirchhausen T, Kozlov MM, Lenz M, Low HH, McMahon H, Merrifield C, Pollard TD, Robinson PJ, Roux A, Schmid S. 2016. Membrane fission by dynamin: what we know and what we need to know. EMBO J 35:2270–2284.

42. AuCoin DP, Smith GB, Meiering CD, Mocarski ES. 2006. Betaherpesvirus-conserved cytomegalovirus tegument protein ppUL32 (pp150) controls cytoplasmic events during virion maturation. J Virol 80:8199–8210.

43. Jones SM, Howell KE, Henley JR, Cao H, McNiven MA. 1998. Role of dynamin in the formation of transport vesicles from the trans-Golgi network. Science 279:573–577.

44. Moorman NJ, Sharon-Friling R, Shenk T, Cristea IM. 2010. A targeted spatial-temporal proteomics approach implicates multiple cellular trafficking pathways in human cytomegalovirus virion maturation. Mol Cell Proteomics 9:851–860.

